# Expert annotation and life-cycle transcriptomics of transcription factors in rust fungi (Pucciniales) highlight the role of cold shock proteins in dormancy exit

**DOI:** 10.1101/2021.10.19.465044

**Authors:** Clémentine Louet, Carla Blot, Ekaterina Shelest, Pamela Guerillot, Jérémy Pétrowski, Pascal Frey, Sébastien Duplessis

## Abstract

Fungi of the order Pucciniales are obligate plant biotrophs causing rust diseases. They exhibit a complex life cycle with the production of up to five spore types, infection of two unrelated hosts and an overwintering stage. Transcription factors (TFs) are key regulators of gene expression in eukaryote cells. In order to better understand genetic programs expressed during major transitions of the rust life cycle, we surveyed the complement of TFs in fungal genomes with an emphasis on Pucciniales. We found that despite their large gene numbers, rust genomes have a reduced repertoire of TFs compared to other fungi. The proportions of C2H2 and Zinc cluster - two of the most represented TF families in fungi-indicate differences in their evolutionary relationships in Pucciniales and other fungal taxa. The cold shock protein (CSP) family showed a striking expansion in Pucciniomycotina with specific duplications in the order Pucciniales. The survey of expression profiles collected by transcriptomics along the life cycle of the poplar rust fungus revealed TF genes related to major biological transitions, e.g. response to environmental cues and host infection. Particularly, poplar rust CSPs were strongly expressed in basidia produced after the overwintering stage suggesting a possible role in dormancy exit. Expression during transition from dormant telia to basidia confirmed the specific expression of the three poplar rust CSP genes. Their heterologous expression in yeast improved cell growth after cold stress exposure, strengthening their implication in dormancy exit. This study addresses for the first time TF involved in developmental transition in the rust life cycle opening perspectives to further explore molecular regulation in the biology of the Pucciniales.

## 1. Introduction

Transcription factors (TFs) are essential for the regulation of expression pathways in eukaryotes by binding genomic DNA in a sequence-specific manner (Charoensawan et al., 2010; Weirauch and Hughes, 2011). TFs are classified according to their DNA-Binding Domains (DBDs) in TF families based on the DBD similarity (Hughes, 2011; Pabo and Sauer, 1992). In fungi, an expansion of TF repertoires was observed in correlation with the genome size, likely explained by a handful of dynamic TF families (i.e. Zn cluster, C2H2 Zn finger and HD-like; Shelest 2008, 2017). Moreover, the comparison across more than 200 fungal genomes determined the presence of approximatively 80 DBD families in fungi, including four fungal-specific families (Shelest, 2017). Several DBD families were associated to regulatory functions of fungal pathogenesis in plant-pathogen interactions, such as sporulation or virulence effector expression (John et al., 2021). Other TFs can be associated to abiotic stress responses; those responsible for temperature stresses are characterized by heat shock (HSDs) or cold shock domains (CSDs). In the latter case, the TFs act as molecular chaperones that destabilize unfavorable secondary structures of mRNAs to facilitate translation after cold shocks (Zhang et al., 2018). CSD proteins (CSPs) are mainly associated with cold adaptation and they function as RNA chaperone with a role in RNA secondary structure stability during cold stress (Chaikam and Karlson, 2010). The structures of CSPs provided evidence for a global conservation of the CSD between prokaryotes and eukaryotes (Chaikam and Karlson, 2010; Heinemann and Roske, 2021) with two consensus RNA-binding motifs (RNP1 and RNP2) mediating RNA binding activity and enhancing the RNA affinity (Landsman, 1992; Nakaminami et al., 2006).

Fungi of the order Pucciniales are major plant pathogens infecting a large diversity of host plants, among which are major crops such as soybean, maize or wheat (Lorrain et al., 2019). The rust diseases caused by these obligate biotrophic pathogens lead to severe loss in agriculture (Figueroa et al., 2018; Godoy et al., 2016). They present singular features such as the capacity to infect one or two alternating hosts (the rust are then termed autoecious or heteroecious respectively) and to form up to five spore types through their complex life cycle (Aime and McTaggart, 2021). Through the year, under temperate conditions, rust fungi can also enter in an overwintering stage during which spores become dormant (Duplessis et al., 2021). In its most complex form, the life cycle of heteroecious rust fungi undergoes a variety of developmental processes that include infection of two taxonomically distant host plants with the formation of specialized infection structures (e.g. germinating tubes, appressoria, infection hyphae, haustoria, spore-forming structures and spores). Moreover, the fungus needs to adapt to the infected tissue and bypass the hosts’ immune systems. These processes require physiological adjustments to nutrient acquisition in the host plants. Sexual and asexual stages also operate on the different hosts and these stages are marked by highly coordinated events. Finally, transitions to dormancy entry and exit are controlled by environmental signals (Duplessis et al., 2021; Mendgen, 1984). Integration of these various signals and developmental processes likely requires a diversity of finely-tuned TF to orchestrate the different genetic programs at play (Duplessis et al., 2021).

In the past ten years, an increasing number of rust genomes have been sequenced (Aime et al., 2017; Lorrain et al., 2019). Most genomes are collected in the JGI Mycocosm database, which allows to draw comparative studies (Grigoriev et al., 2014). Several tools are available to specifically explore the genomes annotations. So far, no global survey of TF genes was conducted in the order Pucciniales, despite reports of gene family expansions presenting typical DBDs (Aime et al., 2017; Cuomo et al., 2017; Duplessis et al., 2011a). Among rust fungi, *Melampsora larici-populina* (poplar rust fungus) is a pioneer model for rust genomics with transcriptomic resources that encompass the whole life-cycle - i.e. infection of the two host plants and the five spore types - making it possible to study specific gene categories based on life-cycle transcriptomics (Duplessis et al., 2021).

In this study, we analyzed the repertoire of transcription factors in Pucciniales compared to other basidiomycetes and to ascomycetes. We showed that despite a striking global contraction of the TF genes complement in Pucciniales, a few families showed significant expansion in comparison to other taxonomical groups, the most remarkable being the CSD family. Life-cycle transcriptomics in the poplar rust fungus showed that CSPs were specifically highly expressed in basidia produced right after the winter dormancy. After confirming these expression profiles in a dormancy interruption experimental setup, we showed that *M. larici-populina* CSP (*MlpCSP*) genes expressed in yeast could improve tolerance to cold stress.

## 2. Material and Methods

### 2.1. Genomic resources and annotation

For genomic analysis, published fungal genomes available in the DoE Joint Genome Institute Mycocosm were used as of the 2021/04/09 (https://mycocosm.jgi.doe.gov/). The TF annotation tables containing PFAM domains were downloaded using the “annotations” track from the JGI Mycocosm for selected taxonomical ranks, i.e. the phyla Ascomycota and Basidiomycota; subphyla Ustilaginomycotina and Pucciniomycotina; and the order Pucciniales. The tables were then filtered from non-TF-like DBDs and non-DBDs. In all comparison analyses, the order Pucciniales and subphyla Ustilaginomycotina and Pucciniomycotina were removed from their respective higher taxonomical ranks, which are thereafter referred to as Other_Pucciniomycotina (i.e. Pucciniomycotina without Pucciniales) and Other_Basidiomycota (i.e. without Ustilaginomycotina and Pucciniomycotina). For each fungal species, the total number of predicted genes and TF-associated domains were collected. CSP sequences were collected as fasta for phylogeny, PCR amplification and cloning (Supplementary Text S1).

### 2.2. Transcriptomic resources

Transcriptomic profiling of *M. larici-populina* was established across its life cycle based on previously published studies transposed to the version 2 of the genome (Guerillot et al., 2021). Briefly, gene expression values were determined for dormant and germinated urediniospores, time course infection of poplar leaves and telia by oligoarrays (Duplessis et al., 2011b; Hacquard et al., 2013) and for basidia, as well as for spermogonia and aecia on larch needles by RNAseq (Lorrain et al., 2018a). Expression values for oligoarrays and RNAseq were normalized separately. For the present study, we extracted expression profiles of *M. larici-populina* TF genes from an open transcriptomic dataset (Supplementary Table S1). The Morpheus tool from the broad institute website (https://software.broadinstitute.org/morpheus/) was used to display normalized transcript expression levels using the *k*-means partitioning method.

### 2.3. Fungal material

Telia from *M. larici-populina* isolate 93GS3 were produced on leaves of the poplar cultivar Robusta (*P. deltoides x P. nigra*) following the procedure described in Pernaci et al. (2014). Briefly, poplar leaves were inoculated by urediniospores at the end of the summer and telia differentiated during autumn in a non-heated greenhouse. Leaves bearing dormant telia were placed outside in natural conditions during winter. Leaves were regularly tested in the lab to ensure that vernalization was reached and then kept at −3°C until further use. Basidia were artificially produced from telia by soaking the leaves in water for six hours at room temperature, followed by 96 hours of incubation at 19 ± 1°C on a wet paper towel in 24 × 24 cm Petri dishes. Pieces of leaves bearing telia (∼20 mg) were sampled right before soaking the leaves in water (t-0h), after 6 hours immersion in water (t-6h) and after 96 hours of incubation (t-96h). For the later condition, both telia which produced basidia visible as a dense orange mat (t-96h_BSD) and non-vernalized telia which did not produce basidia (t-96h_TEL) were separated.

### 2.4. RNA isolation, reverse transcription (RT) and quantitative PCR

Total RNA was isolated from telia and basidia samples with the RNeasy plant mini kit (Qiagen, Courtaboeuf, France), using RLT buffer with ⍰-mercaptoethanol, following manufacturer’s recommendations. The RNase-Free DNase Set (Qiagen) treatment was applied to all samples according to the manufacturer’s recommendations, to eliminate traces of genomic DNA. Primers were designed for *MlpCSP1, MlpCSP2* and *MlpCSP3* (Supplementary Table S2) and efficiency for each target sequence ranged between 80 and 99%. First-strand cDNAs were synthesized from 500 ng DNase-treated total RNA by RT with the iScript cDNA synthesis kit (Bio-Rad) in a total volume of 20 µL according to the manufacturer’s instructions. For the qPCR amplification, 2 µL of one-tenth diluted RT products were amplified in 2X ABsolute qPCR SYBR Green Mix (Thermo Fisher Scientific, Illkirch, France) with 1.6 µM of specific primers with an QuantStudio™ 6 Flex system (Applied, Biosystems, Life Technologies, Saint-Aubin, France) in a final volume of 15µL. Transcript expression levels were normalized with the reference gene *Mlp-ELF1a* using the 2^-ΔCt^ calculation (Livak and Schmittgen, 2001), and expressed relative to the highest expression level, set at 100% (Hacquard et al., 2011b).

### 2.5. cDNA cloning, yeast transformation and cold stress tolerance assay

*MlpCSP1, MlpCSP2* and *MlpCSP3* were amplified from t-96h_BSD cDNA using specific primers defined on the *M. larici-populina* genome sequence (Supplementary Table S2). The PCR-amplified products were digested for 1 h at 37°C with *Bam*HI/*Xho*I and *Hind*III/*Xho*I restriction enzymes for *MlpCSP1*; and *MlpCSP2* and *MlpCSP3*, respectively. Digestion products were purified with the illustra GFX PCR DNA and gel band purification kit (GE Healthcare, Little Chalfont, UK) and were directionally cloned in the yeast expression vector pYES2 (Invitrogen, Thermo Fisher Scientific) carrying the URA3 selection marker driven by the GAL1 promoter; after ligation overnight at 4°C with the T4 DNA ligase (New England Biolabs, Evry, France).The sequences of the cloned fragments in the resulting plasmids pYES2::MlpCSP1, pYES2::MlpCSP2 and pYES2::MlpCSP3 were verified by Sanger sequencing (GENEWIZ, Leipzig, Germany). Yeast strain BY4741 (MATa, Ura3Δ0, Leu2Δ0, His3Δ1, Met15Δ0) grown in YPG medium [1% (w/v) yeast extract, 2% (w/v) peptone, 2% (w/v) glucose] was transformed with each plasmid or with the empty pYES2 plasmid as a control. All transformations were performed according to Gietz and Schiestl (2007). For yeast growth experiments, YNB selective medium [1% (w/v) yeast extract, 2% (w/v) peptone, 2% (w/v) glucose] supplemented with leucine, histidine, methionine was used. Yeast cells transformed with pYES2 constructs and the empty vector were incubated with shaking for 24 hours at 30°C in YNB selective medium as described above. To promote *MlpCSP* expression, cultures were then adjusted to an equal density (OD600 = 0.4) and incubated with shaking for 24 hours at 30°C in YNB inductive medium [1% (w/v) yeast extract, 2% (w/v) peptone, 2% (w/v) galactose] with the amino acids described above. To test the ability of transformed yeast to tolerate cold stress, inducted yeast cells were re-suspended in 2 mL of YNB inductive medium to equal density (OD600 = 1), placed at −20°C during 3, 6, 9, 12 or 24 hours and placed back at 30°C for 12 hours of further growth with shaking (Gao et al., 2012). The relative cell densities were measured after the recovering step at 30°C (OD600). The experiment was performed in triplicate.

### 2.6. Statistical analyses

Statistical analyses were conducted with R software (R Core Team, 2018) for comparisons of means between taxonomical groups based on analysis of variance (Anova) or its non-parametric alternative (Kruskal-Wallis test); followed, when necessary, by appropriate Bonferroni-Dunn’s post hoc test at the threshold level of *P*=0.05.

### 2.7. Phylogenetic analysis of cold-shock proteins

CSP sequences were selected from a collection of representative genomes in distinct taxonomic families available at the Mycocosm for the order Pucciniales (a single representative genome was used for the species *Puccinia graminis* f. sp. *tritici* and *Puccinia striiformis* f. sp. *tritici*, respectively, from Duplessis et al. (2011a) and Cuomo et al. (2017)); a selection of three species from other Pucciniomycotina classes (*Rhodotorula taiwanensis* MD1149 and *Microbotryum lychnidis-dioicae* p1A1 Lamole from the class Microbotryomycetes, and *Mixia osmundae* IAM 14324 v1.0 from the class Mixiomycetes); and the unique CSP in *Aspergilus nidulans* in the phylum Ascomycota as an outgroup. CSP sequences alignment and phylogeny were performed on the Phylogeny website (www.phylogeny.fr) with default parameters in the one click mode (without the Gblock option) combining alignment with Muscle and phylogeny with PhyML (Dereeper et al., 2008). The tree was built with FigTree v1.4.4 (http://tree.bio.ed.ac.uk/software/figtree/) from the exported newick file.

## 3. Results

### 3.1. Annotation of fungal TF domains reveals a contraction of the TF repertoire in the order Pucciniales

In order to have a complete view of the TF category in the order Pucciniales and to compare with other fungi, we took advantage of the updated annotation track at the JGI Mycocosm which compiles a series of 137 PFAMs associated to TF functions. We collected this information for a total of 874 fungal species for which the genomes were published and available for comparative studies (Supplementary Table S3). We performed a curation of this collection of PFAMs based on previous expert annotations (Shelest, 2017, 2008) and ended up with a total of 75 TF PFAMs (Supplementary Table S4). Those excluded lacked a DBD, contained a DBD not associated with a TF function (for instance, those of RNA polymerases), corresponded to general transcription function (e.g. TFII domains) or were related to plant TF with a low representation in the overall dataset. PFAM with a very low representation among all fungi (less than 10 occurrences found among 874 genomes) were also discarded from the set. A total of 58 TF families were defined from their DBD similarity, corresponding to 480339 TF domains in 874 fungal genomes (Table 1).

**Table 1.** Fungal transcription factor families according to their DNA-binding domains (associated PFAM).

| TF gene family name | Associated PFAM (ID: name) |
| --- | --- |
| AFT | PF08731: Transcription factor, AFT |
| AP2 | PF00847: AP2 domain |
| AT-hook | PF02178: AT-hook, DNA-binding motif |
| BRF1 | PF07741: Brf1-like, TBP-binding domain |
| bZIP | PF00170: bZIP transcription factor; PF07716: Basic-leucine zipper (bZIP) region |
| CBFB-NFYA | PF02045: CCAAT-binding transcription factor (CBF-B/NF-YA), subunit B |
| CBFD_NFYB_HMF | PF00808: Histone-like transcription factor (CBF/NF-Y) and archaeal histone |
| Copper-fist | PF00649: Copper fist DNA-binding domain |
| CP2 | PF04516: CP2 transcription factor |
| CSD | PF00313: Cold-shock protein (CSP), DNA-binding |
| DDT | PF02791: DDT domain |
| DP | PF08781: Transcription factor, DP |
| DUF4347 | PF14252: Domain of unknown function (DUF4347) |
| DUF702 | PF05142: Domain of unknown function (DUF702) |
| FAR1 | PF03101: FAR1 DNA-binding domain |
| Forkhead | PF00250: Forkhead domain; PF00498: Forkhead-associated (FHA) domain |
| Fungal-trans | PF04082: Fungal specific transcription factor domain; PF11951: Fungal specific transcription factor domain |
| GCM | PF03615: GCM motif protein |
| GCR1-C | PF12550: Transcriptional activator of glycolytic enzymes |
| HLH | PF00010: Helix-loop-helix DNA-binding domain |
| HMG-box | PF00505: High mobility group (HMG) box; PF09011: High mobility group (HMG)-box domain; PF04769: Mating-type protein MAT alpha 1 HMG-box |
| Homeobox/Homeodomain | PF00046: Homeobox/Homeodomain; PF05920: Homeobox KN domain |
| HSF DNA-bind | PF00447: Heat shock factor (HSF)-type, DNA-binding |
| HTH | PF01381: Helix-turn-helix; PF13560: Helix-turn-helix domain |
| KiIA-N | PF04383: KiIA-N domain |
| Myb | PF00249: Myb-like, DNA-binding domain; PF13837: Myb/SANT-like, DNA-binding domain; PF13921: Myb-like, DNA-binding domain |
| NDT80_PhoG | PF05224: NDT80/PhoG-like, DNA-binding family |
| Not1 | PF04054: CCR4-Not complex component, Not1 |
| NOT2_3_5 | PF04153: NOT2/NOT3/NOT5 family |
| PC4 | PF02229: Transcriptional Coactivator p15 (PC4) |
| PHD | PF00628: PHD-finger; PF13831: PHD-finger |
| PHF5 | PF03660: PHF5-like protein |
| Rap1-DNA-binding | PF09197: Rap1, DNA-binding |
| RFX-DNA-binding | PF02257: RFX DNA-binding domain |
| S1FA | PF04689: DNA-binding protein, S1FA |
| SET | PF00856: SET domain |
| SGT1 | PF07093: SGT1 protein |
| SRF-TF | PF00319: SRF-type transcription factor (DNA-binding and dimerisation domain) |
| STE | PF02200: STE-like transcription factor |
| T-box | PF00907: T-box |
| TBP | PF00352: Transcription factor, TFIID (or TATA-binding protein, TBP) |
| TEA | PF01285: TEA/ATTS domain |
| Tfb2 | PF03849: Transcription factor, Tfb2 |
| TRAUB | PF08164: Apoptosis-antagonizing transcription factor, C-terminal |
| Vhr1 | PF04001: Transcription factor, Vhr1 |
| WRKY | PF03106: WRKY DNA-binding domain |
| YL1 | PF05764: YL1 nuclear protein |
| Zf-C2H2 | PF00096: Zinc finger, C2H2-type; PF12171: Zinc finger, double-stranded RNA-binding; PF13894: C2H2-type zinc finger; PF13912: C2H2-type zinc finger; PF13913: Zinc finger, C2HC-type |
| Zf-C3HC4 | PF00097: Zinc finger, C3HC4-type (RING finger); PF13920: Zinc finger, C3HC4-type (RING finger); PF13923: Zinc finger, C3HC4-type (RING finger) |
| Zf-C5HC2 | PF02928: C5HC2 zinc finger |
| Zf-CCCH | PF00642: Zinc finger C-x8-C-x5-C-x3-H type (and similar) |
| Zf-GATA | PF00320: GATA zinc finger |
| Zf-HC5HC2H | PF13771: PHD-like zinc-binding domain; PF13832: PHD-zinc-finger like domain |
| Zf-met | PF12874: Zinc finger, C2H2-type |
| Zf-NF-X1 | PF01422: NF-X1 type zinc finger |
| Zf-TAZ | PF02135: Zinc finger, TAZ |
| Zn cluster | PF00172: Fungal Zn(2)-Cys(6) binuclear cluster domain |

The order Pucciniales belongs to the subphylum Pucciniomycotina in the phylum Basidiomycota. In order to compare rust fungal TFs, we set taxonomical groups as follows: Pucciniales (n=12), Other_Pucciniomycotina (n=13), Ustilaginomycotina (n=28), Other_Basidiomycota (n=233) and Ascomycota (n=588). In average, the total number of genes is greater in Pucciniales compared with other fungal groups, i.e. 18280 ±1580 in Pucciniales versus 7788 genes ± 500 in Other_Pucciniomycotina, 6945 genes ± 628 in Ustilaginomycotina, 15137 genes ± 687 in Other_Basidiomycota and 10920 genes ± 163 in Ascomycota. On the contrary, whereas the numbers of TF domains and total gene tend to increase concomitantly in all taxonomical groups, the number of TF domains tends to slightly drop in the order Pucciniales (Figure 1). These fungi present a significant reduction relative to Ascomycota and Other_Basidiomycota (Kruskal-wallis, Bonferonni-Dunn test; p-value < 0.0001 and p-value < 0.05, respectively), but not to Other_Pucciniomycotina or Ustilaginomycotina. These results clearly indicate differences between the genome evolution of Pucciniales in comparison to other basidiomycetes and ascomycetes, with a marked contraction in the Pucciniales.

**Figure 1.**
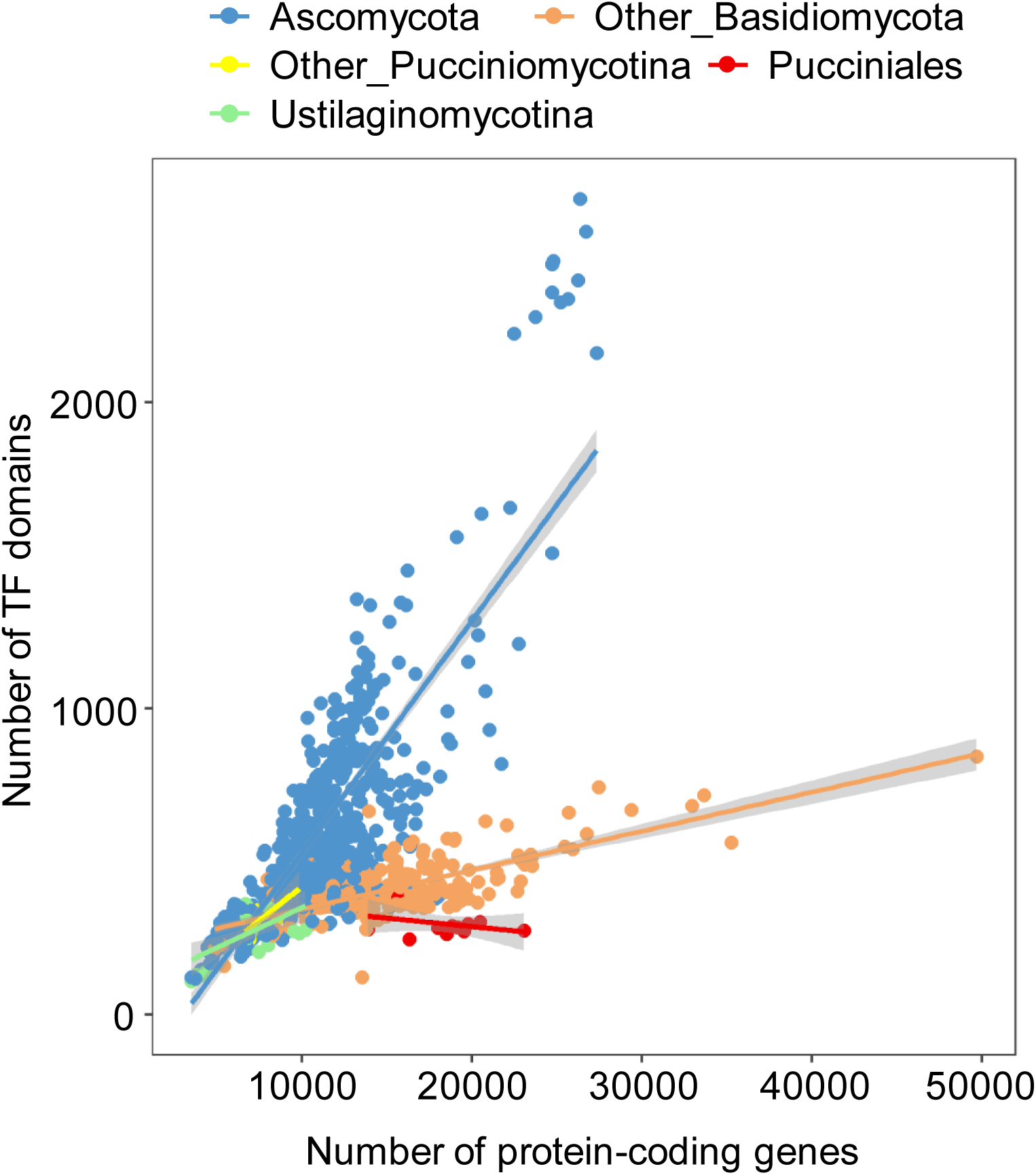
Distribution of the number of TF domains according to the number of protein-coding genes in fungal genomes. Each dot represents a species with a color code according to fungal taxonomical groups. Linear trendlines are indicated for each taxonomical group with the same colour-code as for dots and the level of confidence interval (0.95) is represented in grey. Other_Basidiomycota: Basidiomycota excluding the subphyla Pucciniomycotina and Ustilaginomycotina. Other_Pucciniomycotina: Pucciniomycotina excluding the order Pucciniales.

### 3.2. Comparison of TF families distribution across fungi shows specific evolutionary trajectories in Pucciniales

To evaluate whether the difference observed in fungal TF domains distribution is related to specific contraction or expansion of particular TF families, we extracted the families most represented in the whole dataset (Table 2).

**Table 2.**
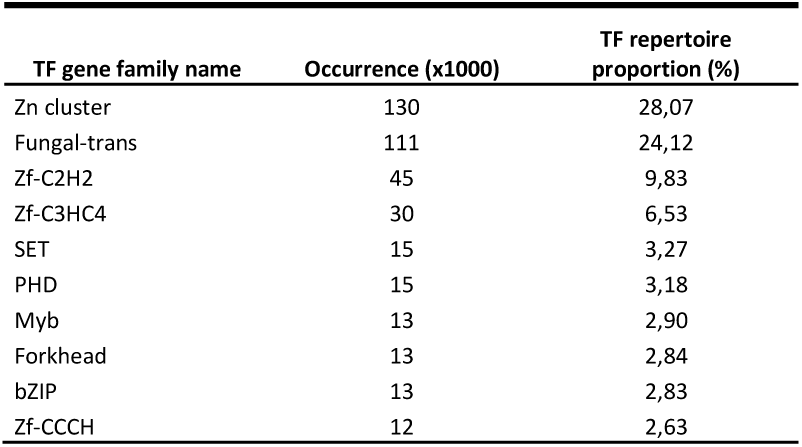
The Top 10 TF families in fungal genomes.

| TF gene family name | Occurrence (x1000) | TF repertoire proportion (%) |
| --- | --- | --- |
| Zn cluster | 130 | 28,07 |
| Fungal-trans | 111 | 24,12 |
| Zf-C2H2 | 45 | 9,83 |
| Zf-C3HC4 | 30 | 6,53 |
| SET | 15 | 3,27 |
| PHD | 15 | 3,18 |
| Myb | 13 | 2,90 |
| Forkhead | 13 | 2,84 |
| bZIP | 13 | 2,83 |
| Zf-CCCH | 12 | 2,63 |

The top 10 TF families (83% of the whole fungal TF domains collection) are shown in Figure 2. The Zn cluster (28.1%), Fungal-trans (24.1%), Zf-C2H2 (9.8%), Zf-C3HC4 (6.5%), and SET (3.3%) families are predominant. Globally, a similar picture can be observed in all surveyed fungal groups. At the contrary, the order Pucciniales presents a different proportion and the four major families are in order Zf-C2H2, Zf-C3HC4, Zn cluster and SET (Figure 2A). Moreover, we can see that whereas the Zn cluster TF dominates in all other fungi, the Zf-C2H2 is more represented in the Pucciniales (ratio of Zf-C2H2/Zn cluster: 1.75 versus 0.62 in average in other fungi; Figure 2B). This clearly points at different evolutionary relationships between the major families of TFs in Pucciniales and other taxa.

**Figure 2.**
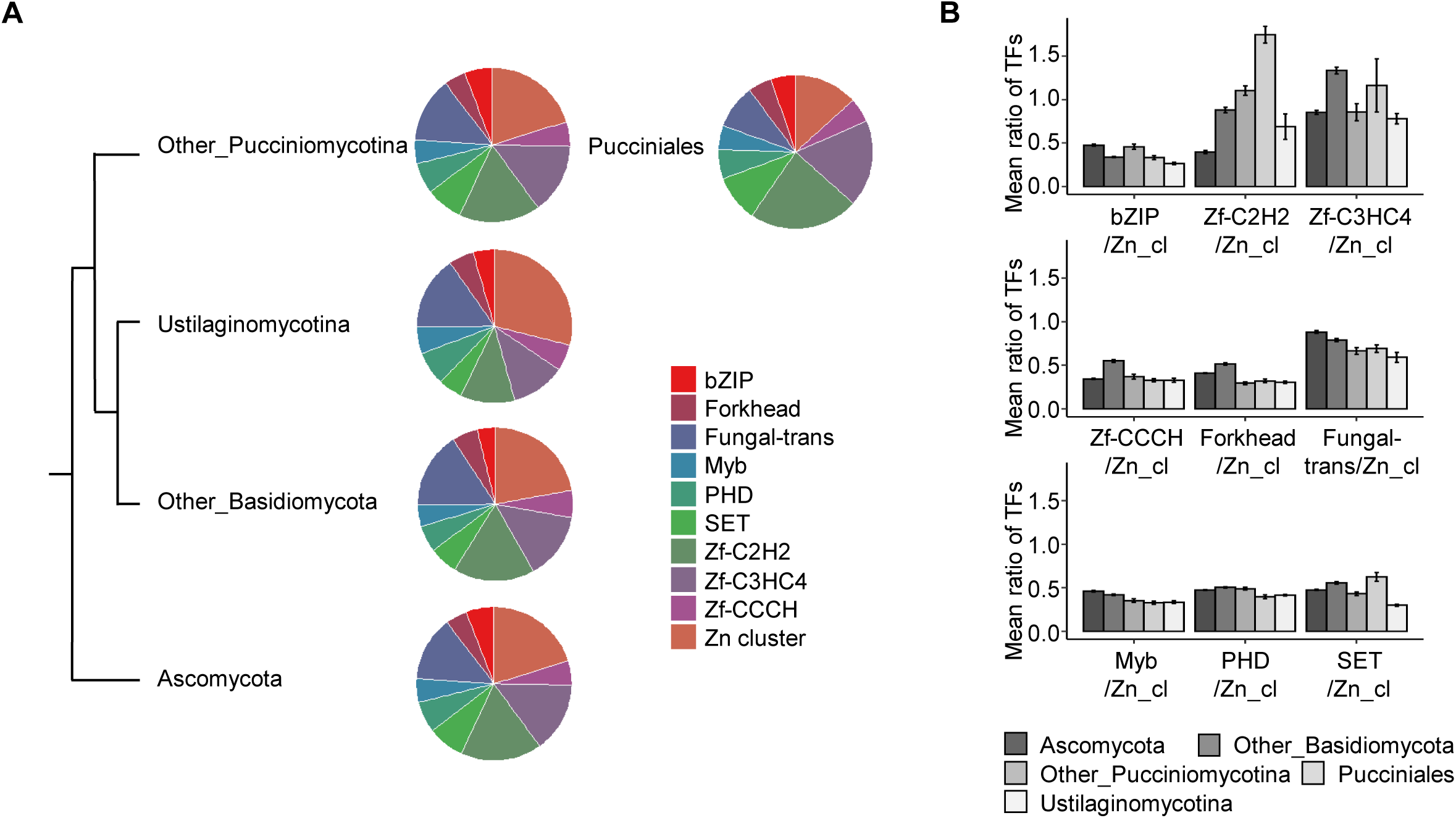
Overview of the proportion of the top 10 TF families in fungal genomes. (A) Pie charts displaying the proportion of the top 10 TF families in fungal taxonomical groups. (B) Ratio comparison of each of the top 10 TF family related to the Zn-cluster family (largest family in Ascomycota and Basidiomycota) for each taxonomical group (average values and standard errors are shown). Other_Basidiomycota: Basidiomycota excluding the subphyla Pucciniomycotina and Ustilaginomycotina. Other_Pucciniomycotina: Pucciniomycotina excluding the order Pucciniales. Tree not to scale according to the fungal phylogeny displayed at the Joint Genome Institute Mycocosm.

### 3.3. Survey of TF families reveals expansion of the CSD family in Pucciniales

To see if there are TF families with unusual distribution, we compared the number of TF domains for each TF family among taxonomical groups. There are 13 families that are significantly differentially represented in Pucciniales than in at least one other group (four over-represented and nine under-represented) and most of them differ from one or two groups (Table 3). The CSD family is outstanding in this respect, being significantly over-represented in the Pucciniales and Other_Pucciniomycotina compared with all other fungi (in average 3.8 and 3.2 CSDs per genome in Other_Pucciniomycotina and Pucciniales, respectively, against 0.07 in all other fungi; Figure 3). Overall, the Pucciniales TF repertoire does not seem to be marked by major expansions or contractions; only the CSD family shows a significant expansion.

**Table 3.** TF families significantly over- or under-represented in Pucciniales related to other fungal taxonomical groups.

| TF gene family name | Comparison | Number of groups | Related taxonomical groups* |
| --- | --- | --- | --- |
| CSD | Over-represented | 3 | Ascomycota (P<0,0001);<br>Other_Basidiomycota (P<0,0001);<br>Ustilaginomycotina (P<0,0001) |
| STE | Over-represented | 2 | Other_Pucciniomycotina (P<0,001);<br>Ustilaginomycotina (P<0,0001) |
| Homeobox/Homeodomain | Over-represented | 1 | Ustilaginomycotina (P<0,01) |
| HSF DNA-bind | Over-represented | 1 | Ascomycota (P<0,0001) |
| PC4 | Under-represented | 3 | Ascomycota (P<0,0001);<br>Other_Basidiomycota (P<0,0001);<br>Other_Pucciniomycotina (P<0,0001) |
| Forkhead | Under-represented | 2 | Ascomycota (P<0,05); Other_Basidiomycota (P<0,0001) |
| Zf-C2H2 | Under-represented | 2 | Ascomycota (P<0,0001);<br>Other_Basidiomycota (P<0,05) |
| bZIP | Under-represented | 1 | Ascomycota (P<0,05) |
| CP2 | Under-represented | 1 | Ascomycota (P<0,0001) |
| HMG | Under-represented | 1 | Other_Basidiomycota (P<0,001) |
| NDT80_PhoG | Under-represented | 1 | Ascomycota (P<0,0001) |
| Zf-CCCH | Under-represented | 1 | Other_Basidiomycota (P<0,05) |
| Zf-GATA | Under-represented | 1 | Other_Basidiomycota (P<0,001) |

\*P-value level after performing a pairwise comparison with a Dunn test (Bonferroni correction)

**Figure 3.**
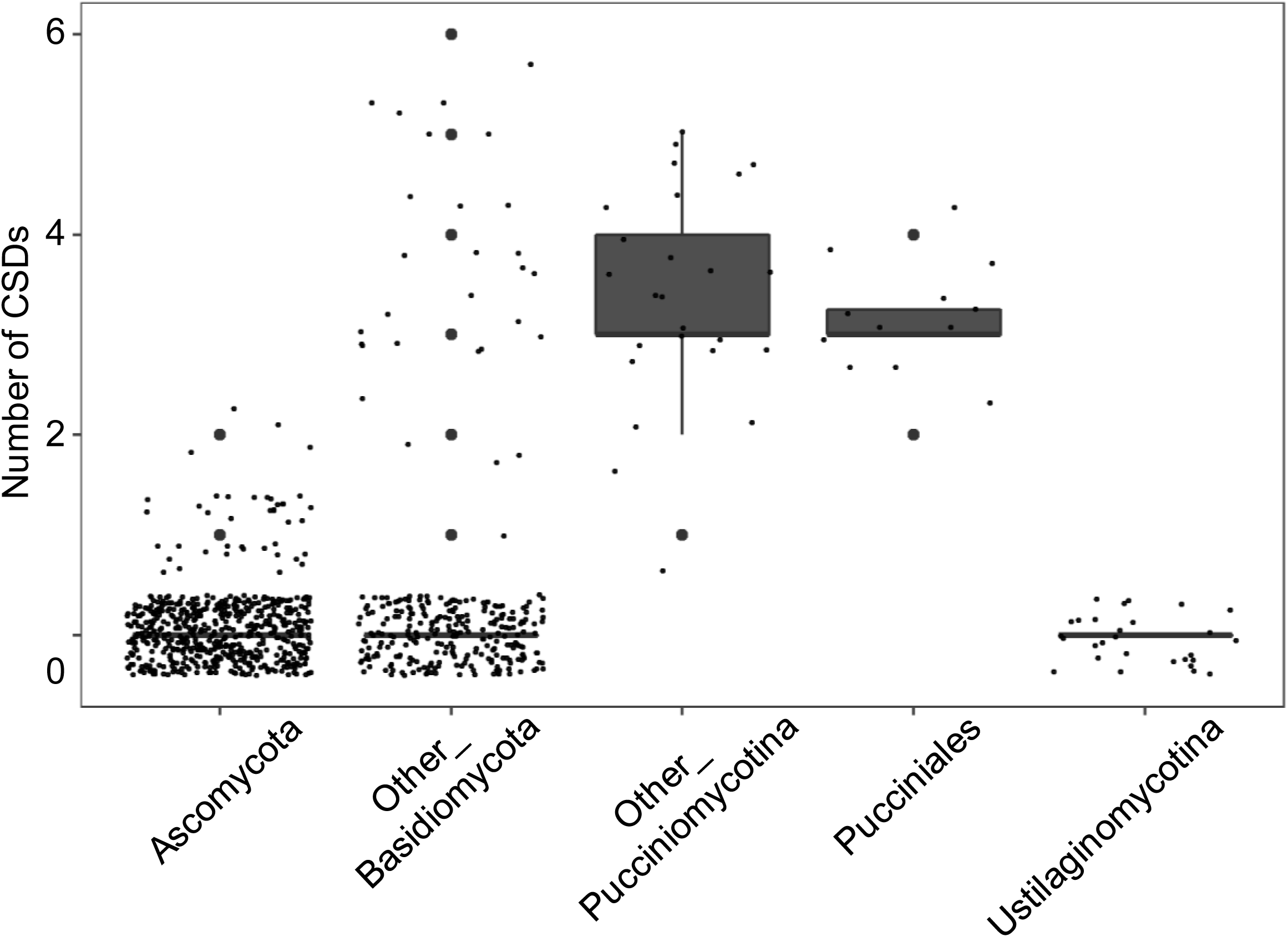
Abundance of cold-shock domains (CSDs) across different fungal taxonomic genomes. Each small dot represents a species. Other_Basidiomycota: Basidiomycota excluding the subphyla Pucciniomycotina and Ustilaginomycotina. Other_Pucciniomycotina: Pucciniomycotina excluding the order Pucciniales. The boxplot outliers are indicated as big dots.

### 3.4. Phylogeny analysis of CSPs supports a specific expansion at the root of the order Pucciniales

To determine the evolutionary relationship between CSPs from the order Pucciniales, we built a phylogenetic tree for CSPs from representative genomes in this order, a selection of three species from other Pucciniomycotina classes, and a single CSP from the phylum Ascomycota (*Aspergilus nidulans*) as an outgroup. The Figure 4 presents the phylogenetic tree (combining alignment with Muscle and phylogeny with PhyML) and a specific focus on the CSD alignment based on 33 full-length amino acid sequences (Supplementary Text S1). The phylogeny showed two major branches for the Pucciniomycotina with a clear dissociation between Pucciniales and other Pucciniomycotina. Two subclades (highlighted in red and green in Figure 4) on one branch and one clade (shown in blue in Figure 4) on the other branch, only consisted of Pucciniales. Within these clades, distinct branches of CSPs belonging to the families Melampsoraceae, Coleosporiaceae and Pucciniaceae were retrieved, supporting the taxonomy of rust fungi. The phylogeny suggests that an ancient duplication occurred at the root of the Pucciniomycotina, and a specific one at the root of the Pucciniales. Furthermore, subsequent duplications occurred independently in two of the three clades for a few rust species (*M. allii-populina, P. triticina*). The CSD alignment showed a relative conservation over the 70 amino acids of the domain compared to the outgroup and the RNP1 and RNP2 motifs exhibit higher levels of conservation (Figure 4). To conclude, this phylogenetic comparison indicates that the CSP expansion likely occurred through ancient and conserved duplications followed by a few more recent ones; with the presence of a conserved CSD containing the RNP1 and RNP2 signature motifs.

**Figure 4.**
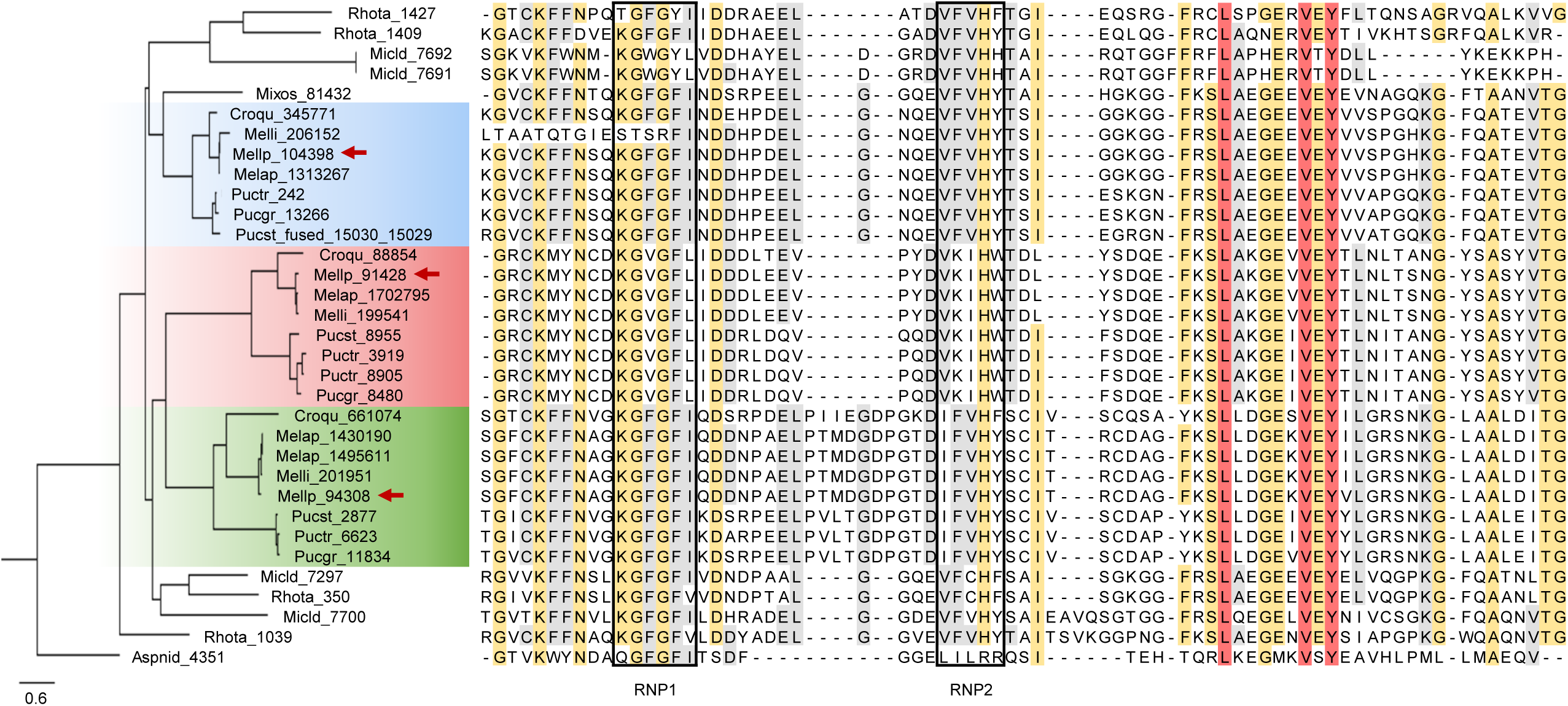
Phylogeny and domain conservation of cold-shock proteins (CSPs) from selected representative genomes in the Pucciniomycotina and the Pucciniales. The phylogenetic tree of 33 full-length fungal CSPs was constructed using the phylogeny.fr website (alignment with Muscle and phylogeny analysis with PhyML) and drawn with FigTree. The unique CSP from the ascomycete *Aspergilus nidulans* was used as an outgroup. The three distinct clades containing Pucciniales CSPs are colored and red arrows indicate the three *Melampsora larici-populina* CSPs. Protein IDs from the Joint Genome Institute Mycocosm are shown on the tree for the different fungal species. The alignment of the cold-shock domain (CSD) is presented next to the tree and amino acid residues are colored according to the level of conservation (red, 100%; yellow, 80%; and light grey, 60%). The RNA-binding motifs RNP1 and RNP2 are indicated by a box according to Prosite.

### 3.5. Transcriptomics of the poplar rust fungus highlights specific expression profiles of TFs during its life cycle

The poplar rust fungus *M. larici-populina* is the only rust fungus for which a survey of the complete life cycle has been established by transcriptomics, covering stages at which all spore forms are formed. Taking advantage of this dataset (Guerillot et al., 2021), we extracted the expression profiles of 213 genes (among a total of 252 TF genes) belonging to TF families. The missing expression profiles correspond to genes for which expression details were not available from the oligoarrays defined on early versions of the genome. The Figure 5 presents the global expression profiling of the 213 *M. larici-populina* TF genes sorted by k-means clustering (see Supplementary Figure S1 for the expanded view of the global profile and *k*-means clusters). Overall, the expression of most genes was not restricted to a single stage, and only a few clusters were predominantly associated with a spore stage (k3, basidia) or an infection stage (i.e. k2 in larch and k4 and k9 in poplar). Expression during time course infection on the poplar host was most often associated with the spore types formed before or after infection (k1, k5, k6, k8 and k10). Interestingly, the k10 cluster corresponds to TF genes expressed at almost all surveyed stages except for early timepoints of poplar infection. Only a limited number of genes showed a specific or preferential expression on a given host, larch (k2; spermogonia and aecia) or poplar (k4, k7 and k9; time course infection), or at a given spore stage (k3; basidia) (see also Supplementary Figure S2). Similarly, only a few TF families could be evidently assigned to a specific stage, such as the CSD family to basidia or the FAR1 family to 24-48 hpi, the earliest infection stages of the poplar time course infection (see Supplementary Figure S1 for profiling according to TF families without grouping). To conclude, this transcriptomic survey allows to derive groups of TF genes coordinately expressed along the poplar rust life cycle and to associate the expression of the FAR1 and CSD families to specific *M. larici-populina* stages. Particularly, the CSD family is only expressed in basidia, a transient stage formed right after the overwintering period.

**Figure 5.**
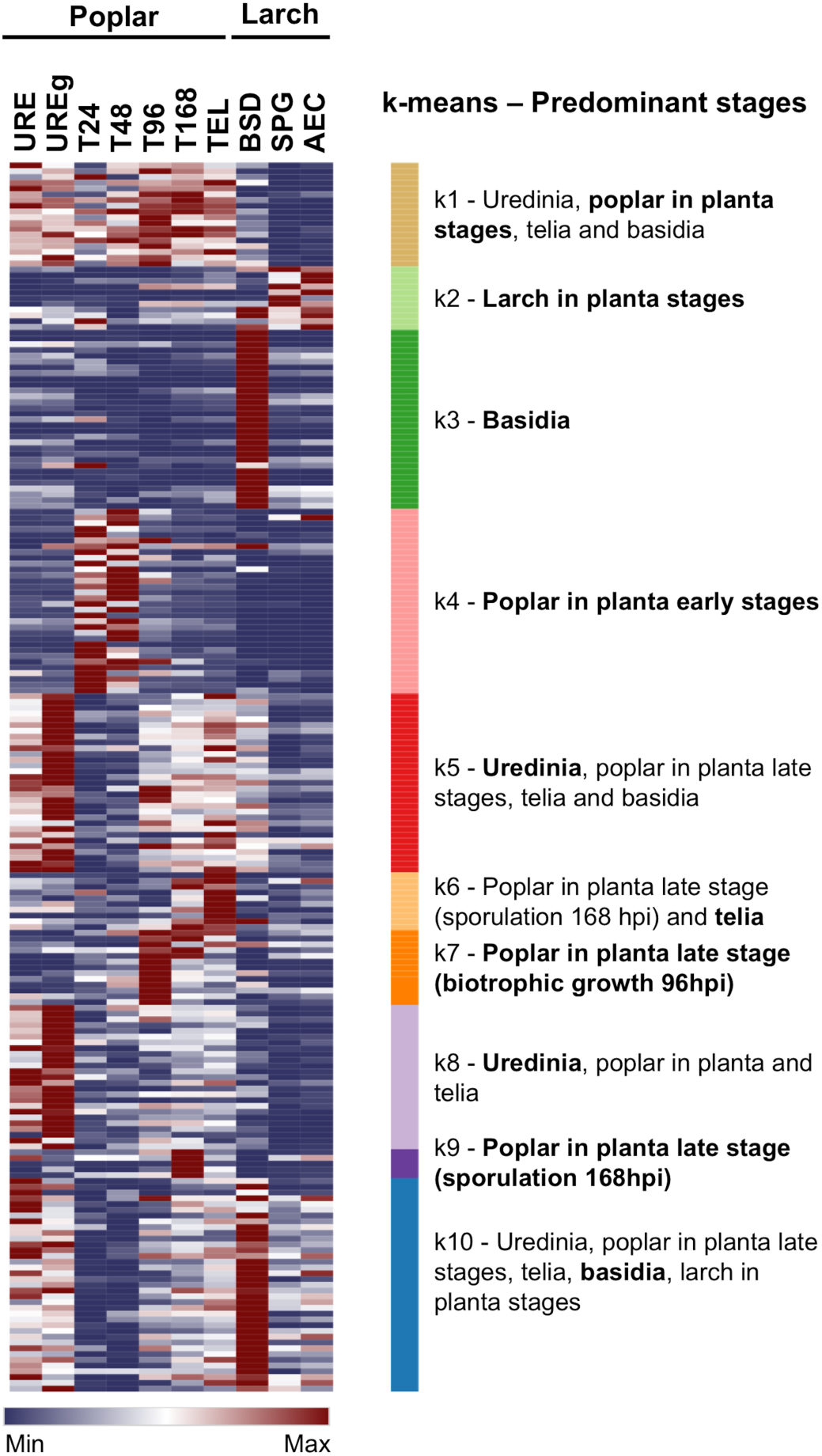
Global expression profiling of the poplar rust fungus TF genes across its life cycle. Expression levels of 213 TF genes were collected from various life stages and grouped by k-means partitioning (k= 10). Life stages are organized according to the yearly poplar rust life cycle starting from stages on the telial host, poplar, followed by stages on the aecial host, larch. Color scale from blue to red represents normalized expression levels from 0 to maximum value at each stage. Biological stages showing high expression levels (i.e. above average values) in each cluster are depicted on the right side of the expression profile and the predominant stage (peak of expression) is highlighted in bold. URE and UREg, dormant and germinating urediniospores, respectively; T24 to T168, hours post-inoculation in a time course infection series on poplar leaves inoculated with urediniospores; TEL, telia; BSD, basidia; SPG, spermogonia; and AEC, aecia.

### 3.6. RTqPCR expression profiling confirms the high induction of poplar rust CSP genes during basidia formation and suggests a role in telia dormancy exit

In order to validate the specific expression of *MlpCSP* genes observed in basidia by RNAseq, we performed a RTqPCR experiment with poplar rust telia and basidia produced in laboratory conditions during the fungal dormancy exit. Similar temporal patterns of transcript expression were observed for the three *MlpCSP* genes (Figure 6) with a relative increase of expression at six hours (telia soaked in water at room temperature) compared with t-0h (dormant telia collected from environmental conditions after an overwintering period). Significantly higher expression levels were then observed for the three genes in basidia produced from overwintered telia after 96 h of incubation at 19°C compared to telia at previous time points (Anova; *p*-value < 0.001 for *MlpCSP1* and *MlpCSP2* and *p*-value < 0.01 for *MlpCSP3*). Expression levels of *MlpCSP* genes in telia that did not produce any basidia after 96 h of incubation (i.e. full extent of dormancy not reached during winter) were not different from the expression in dormant telia. This result validates the specific expression of *MlpCSP* genes in basidia and suggests that the expression could contribute to dormancy exit after the vernalization of poplar rust teliospores.

**Figure 6.**
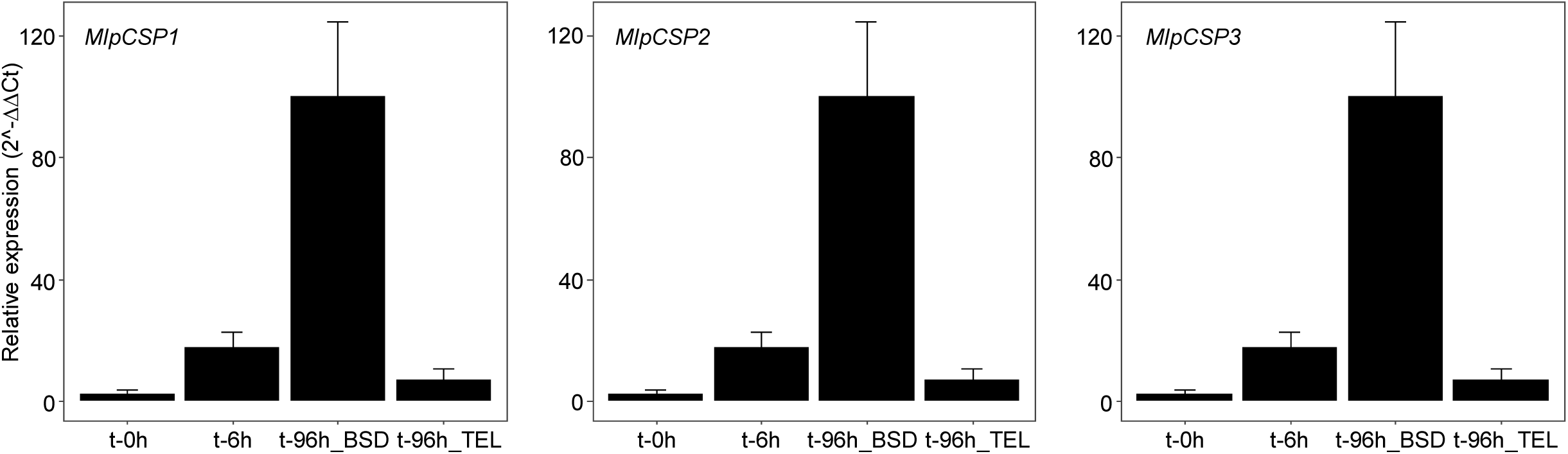
Expression profiles of *Melampsora larici-populina* CSP genes in a dormancy exit experimental setting. Reverse transcription-quantitative polymerase chain reaction (RT-qPCR) was performed for MlpCSP1, MlpCSP2 and MlpCSP3 during a time-course dormancy exit series set on poplar leaves. Expression levels were measured in dormant telia collected from poplar leaves maintained in environmental conditions during winter (t-0h), in telia incubated in water for 6 h at room temperature (t-6h), in basidia differentiated from telia after 96 h at 19°C (t-96h_BSD) and in non-vernalized telia after 96 h at 19°C (t-96h_TEL). Mlp-ELF1a was used as a reference gene for the calculation of each transcript expression levels (2-ΔCt). For each transcript, expression levels are presented as percentages of the highest level measured (2-ΔΔCt). Error bars represent standard deviation of the mean from six biological replicates.

### 3.7. Expression of *MlpCSP* genes in yeast allows cell growth recovery after exposure to cold stress

In the absence of any genetic transformation system for the poplar rust, we expressed each of the three *M. larici-populina* CSP genes in yeast cells to evaluate their functional impact, the yeast genome being deprived of CSP gene. We compared the viability of yeast cells transformed with pYES2::MlpCSP1, pYES2::MlpCSP2 and pYES2::MlpCSP3 plasmids with an empty pYES2 plasmid as a control, after applying different periods of cold stress (i.e. 3 to 24 h of exposure to −20°C) followed by a period of 12 h at an optimal growth temperature of 30°C (Figure 7). Before nine hours of cold treatment, no significant difference could be observed between the control and the different constructs (Anova; *p*-value < 0.05). All transformed yeasts were sensitive to freezing with a decrease of cell viability. From 9 to 24 hours of cold exposure, yeast cells carrying the construct pYES2::MlpCSP2 exhibited a significant higher level of cell growth relative to the control. The yeast cells carrying the pYES2::MlpCSP1 and pYES2::MlpCSP3 constructs showed significant higher levels of cell growth from 12 to 24 hours of exposure to cold compared with the control yeast (Anova; *p*-value < 0.05). Altogether, these results indicate that *MlpCSP1, MlpCSP2* and *MlpCSP3* can all contribute to cold stress tolerance in fungal cells.

**Figure 7.**
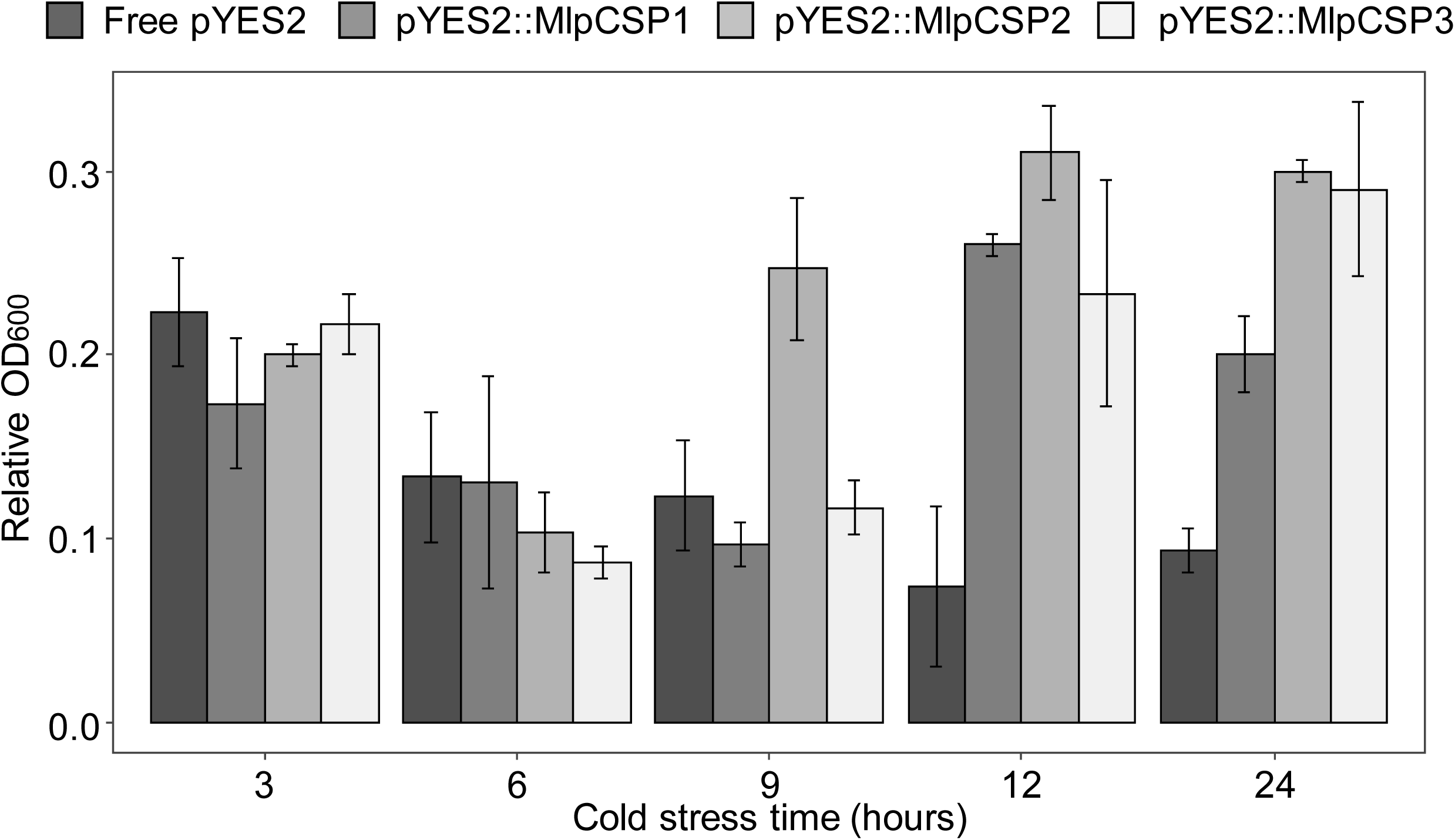
Cell growth recovery assay after cold stress exposure in yeast cells expressing *Melampsora larici-populina* CSP genes. Yeast cells transformed with plasmids pYES2::MlpCSP1, pYES2::MlpCSP2 and pYES2::MlpCSP3 and empty pYES2 as a control were incubated in an inductive medium for 24 hours at 30°C. The ability of transformed yeast cells to tolerate cold stress was tested by placing the cells at −20°C during 3, 6, 9, 12 or 24 hours and back at 30°C for 12 hours of further growth. The relative cell densities were measured after the recovering step at 30°C (OD600) and compared to cell densities before freezing. Values correspond to the mean of three biological replicates and error bars indicate the standard deviation of the mean.

## 4. Discussion

Rust fungi are obligate plant pathogens with a unique biological cycle among eukaryotes marked by an alternation of up to five different spore stages differentiated on one or two different host plants (Lorrain et al., 2019). TFs are key cellular elements to finely tune the regulation of genetic programs underlying developmental transitions influenced by both host and environmental signals (Duplessis et al., 2021). Studies of TFs in rust fungi are scarce. The homolog of *S. cerevisiae* STE12 involved in morphological transitions and sexual development was studied in the wheat stripe rust fungus *P. striiformis* f. sp. *tritici*, in which *PstSTE12* silencing inhibits the pathogen growth *in planta* (Zhu et al., 2018). Also, the homeodomain pair HD1 and HD2 which play a critical role during mating in fungi were cloned from the wheat rust *Puccinia triticina* and were proven to be active when expressed in the heterologous system *Ustilago maydis* (Cuomo et al. 2017). So far, apart from global surveys in fungi (Shelest, 2017, 2008), no dedicated analysis of TF was conducted in rust fungi. Here, by combining a dedicated annotation of TF genes and transcriptomics in a model rust fungus, we address some biological specificities in rust fungi related to their peculiar life cycle.

A previous survey of the TFome in fungal genomes showed that TF numbers tend to increase with the size of the gene complement (Shelest, 2017). Strikingly, whereas rust fungal genomes present significantly larger gene numbers than other fungi, they present reduced TF repertoires. On the other hand, some TF families have had peculiar evolutionary history. For instance, the proportion of Zf-C2H2 related to Zn cluster is higher in Pucciniales and Pucciniomycotina when compared to other fungi. The Zn cluster is the most abundant TF family in fungi and more particularly in younger Ascomycota taxa; and they are associated to a variety of regulatory processes (Shelest, 2017; Weirauch and Hughes, 2011). A future challenge will be to determine which specific Zn cluster and Zf-C2H2 TF genes have been retained or lost in Pucciniales and whether their regulatory roles and target genes are conserved. Apart from these striking differences, only a handful of gene family expansions or contractions could be observed, with CSPs standing out in this respect. Whereas CSPs are absent from a large majority of ascomycetes and basidiomycetes, they are significantly expanded in the order Pucciniales and in other Pucciniomycotina, which exhibit up to five genes. As our analysis suggests, CSP expansion originates from successive duplications in the subphylum Pucciniomycotina and three types of CSP genes belonging to the same branch can be distinguished in the phylogeny of the order Pucciniales. Fungi of the Pucciniomycotina have been reported in a diversity of ecological habitats, including extreme environments imposing strong abiotic stresses (e.g. artic ice, deep-sea habitats; Aime et al., 2014; Toome-Heller, 2016). Pucciniales that are found in temperate, montane or boreal climates most often display a dormant phase during winter in their cycle (Duplessis et al., 2021). The expansion of CSP genes in Pucciniomycotina suggests a role in their adaptation to cold environment.

Transcriptomics is an approach of choice to decipher TF functions in eukaryotes. By combining different transcriptomic studies conducted in the model rust fungus *M. larici-populina* (Guerillot et al., 2021), it is possible to specifically survey TF expression along the rust life cycle in order to infer TF functions. As expected for this cellular category, a variety of TF genes from different families are expressed at each life stage. Specific TFs associated with a given stage or with a given host plant can be identified. Interestingly, nearly 80% of the TFs with the highest expression levels were previously reported as regulators of key virulence pathways in phytopathogenic fungi (John et al., 2021). During time course infection of the poplar host, preferential expression of TF genes can also be associated with early colonization, biotrophic growth or sporulation. Similar profiles were previously described for other categories such as virulence effectors (Duplessis et al., 2011b; Hacquard et al., 2012), and it will be crucial to determine whether specific TF can be assigned to such regulation. Moreover, the expression of only few TF families could be restricted to a given stage. For instance, almost all genes of the FAR1 family show a specific expression at the earliest stages of poplar infection, whereas they are not detected in sporulating spores. At this stage, infection hyphae are colonizing the host mesophyll (Hacquard et al., 2011a). An assumed role of FAR1 is mobilization of intracellular lipid reserves during plant infection by the phytopathogenic fungus *Magnaporthe oryzae* (bin Yusof et al., 2014). It would be interesting to decipher the exact role of FAR1 at this stage and whether regulation of lipid metabolism also plays a role at this stage of rust infection.

*M. larici-populina* CSPs exhibit the most remarkable expression profile among TF genes, restricted to the basidia stage with the highest transcript levels across the global expression dataset. Basidia are produced from telia right after overwintering when favorable conditions are met (Duplessis et al., 2021). This stage is transient and basidiospores rapidly differentiate to realize infection on larch, the alternate host of the poplar rust fungus (Hacquard et al., 2011a; Pernaci et al., 2014). Temperature and moisture seem to be crucial factors to exit winter dormancy (Mendgen, 1984). An experimental set-up mimicking dormancy exit confirmed the high expression of the three CSP genes, highlighting their major role in the transition from cold to favorable temperatures. The biotrophic status of rust fungi impedes their genetic manipulation and gene functional characterization requires heterologous systems (Bakkeren and Szabo, 2020; Hu et al., 2007; Lorrain et al., 2018b). In order to investigate the role of CSP function, the three *MlpCSPs* were expressed in yeast cells to assay cold recovery as previously described for other fungi (Fang and Leger, 2010). Yeast cells expressing the different rust CSPs showed an improved growth after exposure at −20°C compared to control yeast devoid of CSP. Here, we show that the three *MlpCSPs* are highly expressed in basidia and can improve growth of yeast cells after a cold shock. Altogether our results clearly indicate that CSPs are major TFs of the rust life cycle with a role associated to dormancy exit and basidia differentiation. In other eukaryotes a variety of functions have been assigned to CSPs (Heinemann and Roske, 2021). A next challenge will be to determine whether each of the three *MlpCSP* genes exhibit some specificity or show functional redundancy in the regulation of subsequent cellular processes.

To conclude, by combining functional annotation in rust genomes and life cycle survey in transcriptomics data, we open the way to decipher key regulatory processes in the complex biology of rust fungi. Most often, only infection on the main host is studied in rust fungi precluding the understanding of other stages. Our strategy allows to address for the first time the function of regulatory genes in basidia of rust fungi. Beyond the case of CSPs, other major categories may now be considered for detailed analysis at different stages. Fine dissection of specific rust stages by omics approach should help to better define the genetic programs correlating with TF expression.

## CRediT authorship contribution statement

**Clémentine Louet:** Conceptualization, Methodology, Validation, Formal analysis, Investigation, Resources, Data Curation, Writing - Original Draft, Writing - Review & Editing, Visualization, Supervision. **Carla Blot:** Validation, Formal analysis, Investigation, Resources, Data Curation. **Ekaterina Shelest:** Methodology, Formal analysis, Investigation, Resources, Data Curation, Writing - Review & Editing. **Pamela Guerillot:** Methodology, Software, Validation, Formal analysis, Investigation, Resources, Data Curation, Writing - Review & Editing. **Jérémy Pétrowski:** Resources. **Pascal Frey:** Resources, Writing - Review & Editing, Funding acquisition. **Sébastien Duplessis:** Conceptualization, Methodology, Software, Validation, Writing - Original Draft, Writing - Review & Editing, Visualization, Supervision, Project administration, Funding acquisition.

## Acknowledgements

We would like to thank the Redox team of the UMR IAM for hosting molecular experiments, providing inputs and material as well as for rich discussions along the way. We would like to thank all colleagues who were involved in generating primary poplar rust transcriptomic data that we exploited here.

## Appendix A. Supplementary material

**Supplementary Table S1**. Expression values of the poplar rust fungus TF genes.

**Supplementary Table S2**. Primers for cloning and expression analysis of *Melampsora larici-populina* cold shock domain (MlpCSP) genes.

**Supplementary Table S3**. List of fungal genomes from JGI.

**Supplementary Table S4**. List and occurrence of TFs detected in fungal genomes.

**Supplementary Figure S1**. Expression profiling of *Melampsora larici-populina* TF genes grouped by k-means (k= 10) across the life cycle. **A**. Global expression profile with genes sorted by TF families. **B**. Global expression profile with genes sorted by k-means. Expression values scale from blue to red represents normalized expression levels from 0 (deep blue) to maximum value (deep red) at each stage. Morpheus (https://software.broadinstitute.org/morpheus) parameters used for k-means were as follows: one minus pearson’s correlation, cluster by rows, number of clusters 10 (1000 iterations). For each gene, the id shown on the right side of the profile indicates the Joint Genome Institute Mycocosm protein-ID number with the associated PFAM and TF family in brackets. The 10 k-means are shown on the right side of each profile with k-means colors corresponding in A and B.

**Supplementary Figure S2**. Top 10 most highly expressed *Melampsora larici-populina* TF genes at each stage of the poplar rust life cycle. The life cycle is shown ordered from stages occurring on the telial host to those occurring on the aecial host. URE and UREg, dormant and germinating urediniospores, respectively; T24 to T168, hours post-inoculation in a time course infection series on poplar leaves inoculated with urediniospores; TEL, telia; BSD, basidia; SPG, spermogonia; and AEC, aecia. The top 10 most highly expressed TFs have been grouped according to the life cycle as stage-specific, host-specific or other (not host- or stage-specific) based on the stage where they showed highest expression levels.

**Supplementary Text S1**. List of cold-shock amino-acid sequences used in this study.

