## Supplementary figures and images for "Expert annotation and life-cycle transcriptomics of transcription factors in rust fungi (Pucciniales) highlight the role of cold shock proteins in dormancy exit"

### Supplementary Figure S1

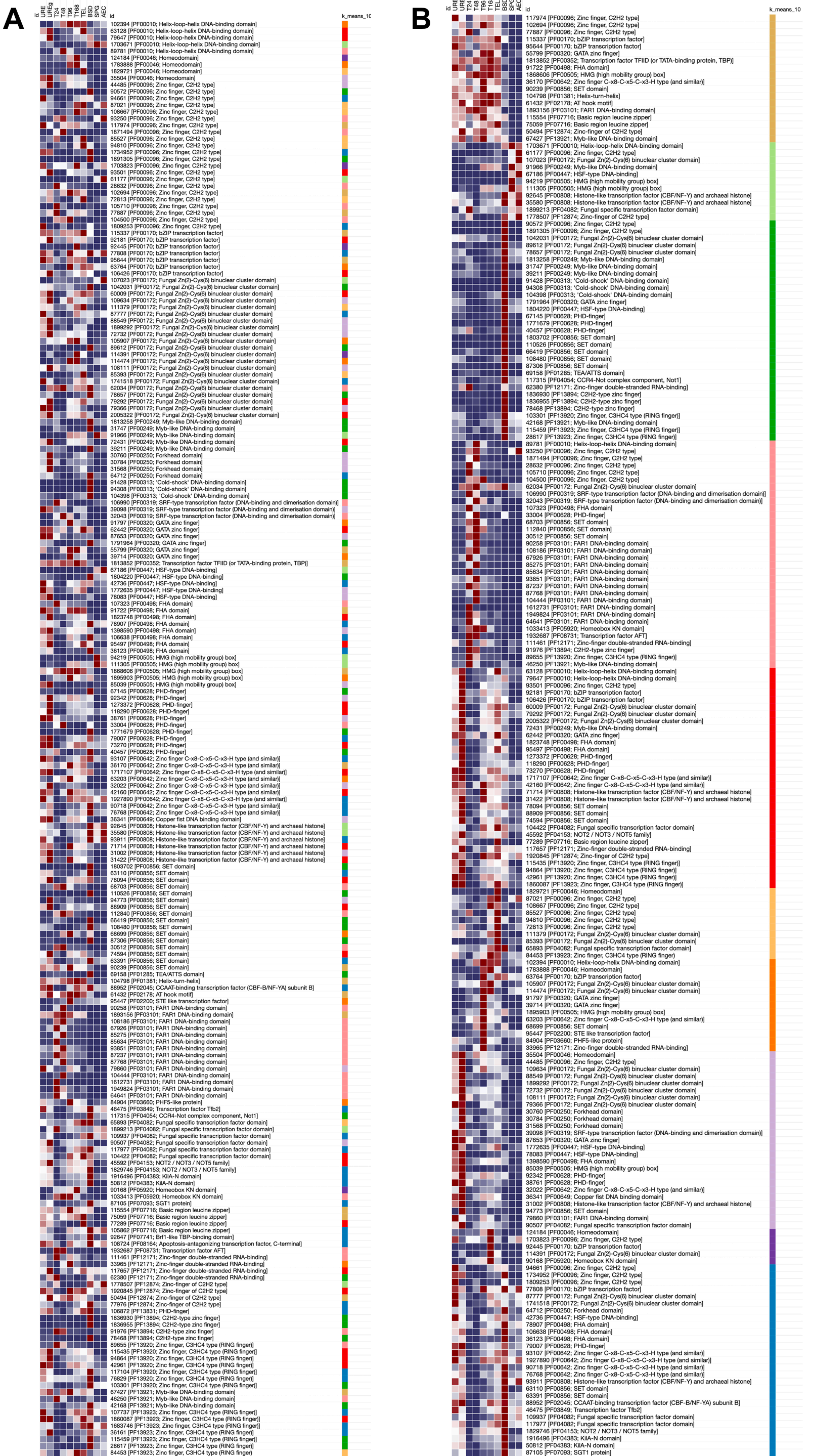

### Supplementary Figure S2

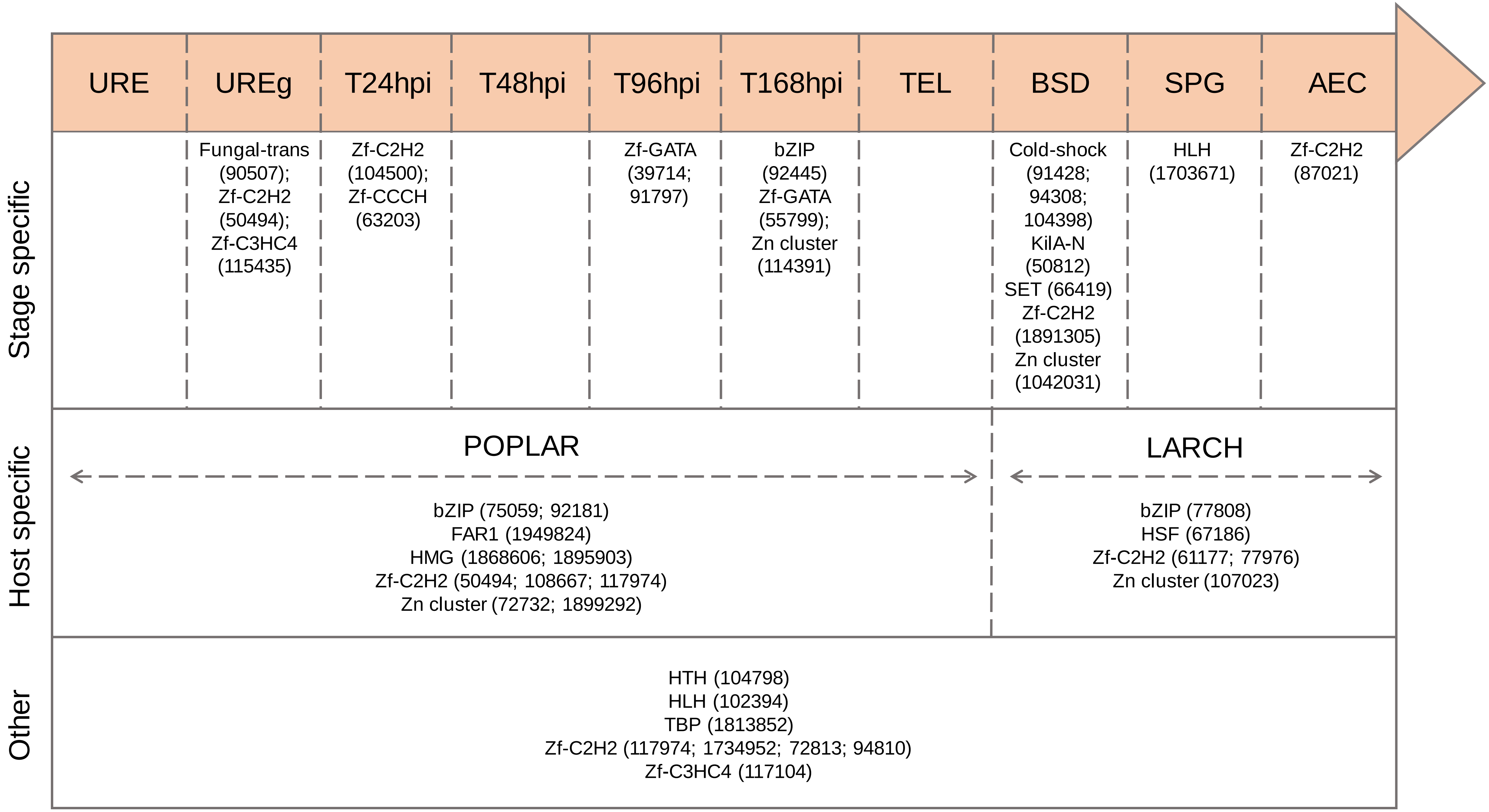
